# Rfpred: A Random Forest Approach for Prediction of Missense Variants in Human Exome

**DOI:** 10.1101/037127

**Authors:** Fabienne Jabot-Hanin, Hugo Varet, Frederic Tores, Alexandre Alcaïs, Jean-Philippe Jaïs

## Abstract

Exome sequencing is becoming a standard tool for gene mapping of genetic diseases. Given the vast amount of data generated by Next Generation Sequencing techniques, identification of disease causal variants is like finding a needle in a haystack. The impact assessment and the prioritization of potential pathogenic variants are expected to reduce work in biological validation, which is long and costly.

One of the possible approaches to determine the most probable deleterious variants in individual exomes is to use protein function alteration prediction. We propose in this paper to use a machine learning approach, the random forest to build a new meta-score based on five previously described scores (SIFT, Polyphen2, LRT, PhyloP and MutationTaster) and compiled in the dbNSFP database.

The functional meta-score was trained on a dataset of 61 500 non-synonymous Single Nucleotide Polymorphisms (SNPs). The random forest method (rfPred) appears to be globally better than each of the classifiers separately or in combination in a logistic regression model, and better than a newly described score (CADD) on independent validation sets.

RfPred scores have been pre-calculated for all the possible non-synonymous SNPs of human exome and are freely accessible at the web-server http://www.sbim.fr/rfPred/

## 1. Introduction

Exome sequencing is a recent and important innovation for the exploration of patients affected by a genetic disease, in particular in situations when the other approaches have failed. The difficulty is often to find the disease causal variant(s), and it may be useful to focus on computational approaches like functional alteration predictors, to synthesize all available a priori information for variants prioritization before further functional studies.

Many different methods have been developed and published over the past fifteen years, each of these has distinct advantages and disadvantages, but none can be considered as the gold standard ^1^^2^^3^. The prediction scores of some of these methods have been compiled in the dbNSFP database for all known protein coding genome positions ^4^. Besides, Li and colleagues proposed to combine five of them in a logistic regression framework ^5^ in order to globally improve predictive performance in comparison with individual scores.

## 2. Materials and Methods

### 2.1 *Data collection for model building*

First, we constructed a Single Nucleotide Variant (SNV) dataset with a status variable, taking values “neutral” or “deleterious”, in order to build the prediction model. For the deleterious variants, we used the OMIM database (Online Mendelian Inheritance in Man) ^7^ - 23/09/2001 version- available at https://main.g2.bx.psu.edu/library, which inventories variants and phenotypes associated with Mendelian diseases. It contains 9130 human genome variants using the hg19 map. These variants will be considered as deleterious.

To build an assumed neutral variant set, we started with the 1000 genome database available via ANNOVAR ^8^ http://www.openbioinformatics.org/annovar/ (version from November 2010 updated in June 2011), which inventories the genetic data of supposed healthy subjects. Among these data, we selected the missense variants with the already existing hg19_avsift filter of ANNOVAR, and those with an allele frequency < 1% in the population (rare variants); their neutral nature is not obvious and corresponds to the reality with which researchers are faced.

For each of these variants, a score has been attributed with the five following methods: SIFT (released August 2011) ^9^, Polyphen2 (HumDiv classifier model v2.1.0) ^10^, Mutation Taster (released March 2010) ^11^, LRT (released November 2009) ^12^ and PhyloP ^13^ thanks to the dbNSFP public database https://sites.google.com/site/jpopgen/dbNSFP. ^4^. This database contains all possible SNPs within human genome coding regions, which have been determined by the CCDS project ^14^, and for each of the 87 million SNPs, the scores of the five predictors have been precalculated and made available. The scores are used raw (directly calculated by the softwares) or processed such that the pathogenicity probability increases with increasing score.

We have kept in this training dataset only variants with the five available scores, which can be reduced to 6 254 deleterious SNPs and 55 223 neutral SNPs.

### *2.2* *Data Collection for external validation*

In order to evaluate the prediction model on independent datasets, we have used 2 general and 2 more precise datasets:

- A published variant dataset – EXOVAR – already used to evaluate similar methods (available at http://statgenpro.psychiatry.hku.hk/limx/kggseq/download/ExoVar.xls) including both 4752 neutral (from 1000 Genomes Project^15^, with a derived allele frequency <1%) and 5340 disease causing variations (with known effects on the molecular function causing human Mendelian diseases from the UniProt database) ^16^, but only 1740 neutral and 3601 disease causing variants not included in our learning dataset with all necessary scores available.
- A validation dataset composed of 1100 pathogenic non-synonymous variations coming from ClinVar database (annotated as “Pathogenic”) and of 5412 variations coming from 1000 Genomes Project^15^ with a minor allele frequency between 5% and 20%, considered as non-deleterious and for which the 5 predictors give a score. None of these variations are part of our learning dataset.

Two more specific missense genetic variant datasets coming from two genes having many deleterious or polymorphic known variants:

- The first one is the *COL4A5* gene on the X chromosome which is composed of 51 exons making up a total length of 257kb. Many variants in this gene are implied in the Alport syndrome [MIM 301050], but SNPs without known disease association are also reported in public databases. After having discarded the SNPs included in our learning dataset, this first validation dataset contains:

- 34 neutral variants coming from the dbSNP database – build 137 ^17^, with the “clinical significance” annotation in the database different from “probable pathogenic”
- 168 deleterious variants coming from dbSNP database - build 137 also, annotated “probable pathogenic”, and from HGMD database (public version) ^18^.
- The second analyzed gene is the *COL7A1* gene on the chromosome 3 which is composed of 118 exons making up a total length of 32kb. Some variants in this gene are associated in the dystrophic epidermolysis bullosa [MIM 131750]. Our validation dataset contains:

- 325 neutral variants coming from the dbSNP database – build 137, with the “clinical significance” annotation in the database different from “probable pathogenic”
- 162 deleterious variants coming from dbSNP database – build 137 annotated “probable pathogenic” and the col7info database ^19^.

### *2.3* *Statistical method*

Relationships between the five scores on the training dataset were analyzed by computing the Spearman’s rank correlation matrix. The meta-score rfPred was derived from the five individual scores using the predictions of a random forest model^20^.

Briefly a random forest is based on a set of classification trees trained on a random subset of observations and variables of the complete dataset. Each tree votes for a given class, here either neutral or deleterious variant, and the final classification is based on the trees vote’s majority. In fact, more precise information is provided by the proportion of tree votes in favor of the deleterious class and it can be used as a credibility index of the classification. So we used this vote proportion as a predictive meta-score of functional alteration.

We generated a random forest model composed of 5 000 trees based on two random scores among the five available for splitting at each tree node, using the model deviance as classification criterion. Each tree was trained against an equal sampling of 2000 variants of each class (deleterious and neutral).

All the analyses were made with R software version 2.15.0, and in particular with the package named “randomForest” ^21^ ^22^.

### *2.4* *Statistical model evaluation*

To compare the predictors’ performances with the rfPred one, we have used the individual prediction scores (Polyphen2, SIFT, LRT, PhyloP and MutationTaster) coming from dbNSFPv2.0 ^23^ and CADD scores computed via its web-interface^6^. We have then computed the ROC curves and the areas under the curves of the different classifiers thanks to the R package “Hmisc”^24^ on the learning dataset and on the validation datasets.

In order to have a more useful comparison between the classifiers, we have considered also two particular scenarii corresponding to the reality faced by molecular biologists:

In the one hand, the classification model is used on a very long variants list coming directly from the exome alignment on the reference genome in order to establish a list a potentially pathogen variants. The purpose in this case is to keep all potentially deleterious variants and not to exclude any potential candidate. We favor in this case the sensitivity to the detriment of the specificity.

In the other hand, a variant selection process has been made beforehand by other methods (typically existing methods in pipelines of next-generation sequencing platforms and/or prior knowledge about the genetic basis of the disease), and the classification model will be used in a second time to prioritize the work areas. The goal is to minimize false positives variant rates to concentrate biological efforts on variants with a high probability of pathogenicity. We thereby favor specificity.

For these specific scenarii, partial AUC (Areas Under Curve) restricted to False Negative Rates < 10% for the first scenario and to False Positive Rates < 10% for the second one, as well as the ratio 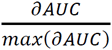 (partial area index) have been calculated.

MutationTaster has not been evaluated on the ClinVar-1000Genomes dataset because a very large part of the variants have been used to train the method.

### *2.5* *Missing data handling*

For some exome positions, one or several pre-calculated scores were missing in dbNSFP 2.0 ^23^. Because we wanted to be able to use rfPred even if one of the score is missing for a variant position, we have imputed the missing score value by a random forest approach implemented in the R package “yaImpute” ^25^, based on a k-NN algorithm. We have used k=1 in our procedure.

## 3. Results

Spearman’s correlation matrix for the individual predictors is given in Table 1. It shows low to moderate correlation between scores (0.18 for the minimal and 0.66 for the maximal correlation) on the learning dataset, indicating that the information contained in the five prediction scores should not be completely redundant and that a combination of them could therefore be pertinent.

**Table 1:** Spearman correlation coefficients matrices on learning dataset with OMIM deleterious variants and neutral variants with allele frequençy < 1%

| Datasets |  | PhyloP | SIFT | Polyphen2 | LRT | MutationTaster |
| --- | --- | --- | --- | --- | --- | --- |
| Neutral Variants<br>From 1000 genomes<br>(Allele frequency < 1%) | PhyloP | 1 | 0.28 | 0.43 | 0.6 | 0.47 |
|  | SIFT |  | 1 | 0.58 | 0.28 | 0.32 |
|  | Polyphen2 |  |  | 1 | 0.45 | 0.47 |
|  | LRT |  |  |  | 1 | 0.66 |
|  | MutationTaster |  |  |  |  | 1 |
| Deleterious Variants<br>From OMIM | PhyloP | 1 | 0.18 | 0.22 | 0.35 | 0.23 |
|  | SIFT |  | 1 | 0.52 | 0.33 | 0.38 |
|  | Polyphen2 |  |  | 1 | 0.43 | 0.42 |
|  | LRT |  |  |  | 1 | 0.53 |
|  | MutationTaster |  |  |  |  | 1 |

### *3.1* *Model prediction performance*

The random forest prediction model (rfPred) has an AUC of 0,849 on the “Out of bag” learning dataset (unbiased AUC), whereas the best individual predictor on the same dataset, MutationTaster, obtains only 0,804 of AUC. The simple logistic model meanwhile obtains a maximal AUC of 0,827, even if the AUC is computed on the totality of the sample used to build the model. About CADD, it reaches the AUC of 0,770. We can note that rfPred ROC curve lies above all the others on the whole curve. (Figure 1)

**Fig 1:**
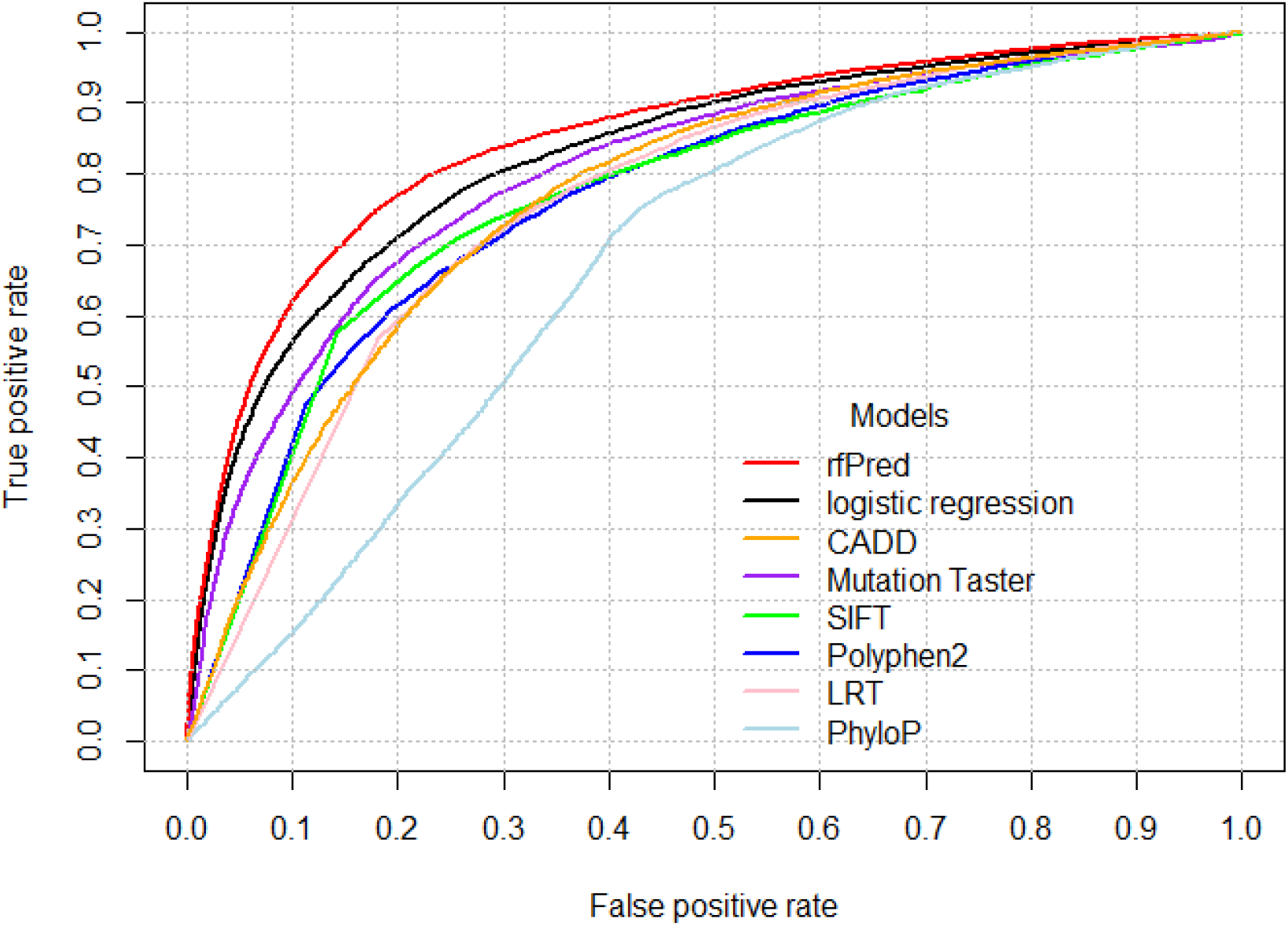
Classifiers’ ROC curves on learning dataset

The variable importance is measured in the rfPred model by the mean decrease in Gini coefficient.

It is a measure of how each variable contributes to the homogeneity of the nodes and leaves in the resulting random forest. Each time a particular variable is used to split a node, the Gini coefficient for the child nodes are calculated and compared to that of the original node. The Gini coefficient is a measure of homogeneity from 0 (homogeneous) to 1 (heterogeneous). The changes in Gini are summed for each variable and normalized at the end of the calculation. Variables that result in nodes with higher purity have a higher decrease in Gini coefficient.

Table 2 indicates that the two main predictors are Mutation Taster and SIFT scores for rfPred model. This is consistent with the layout of the observed ROC curves.

**Table 2:**
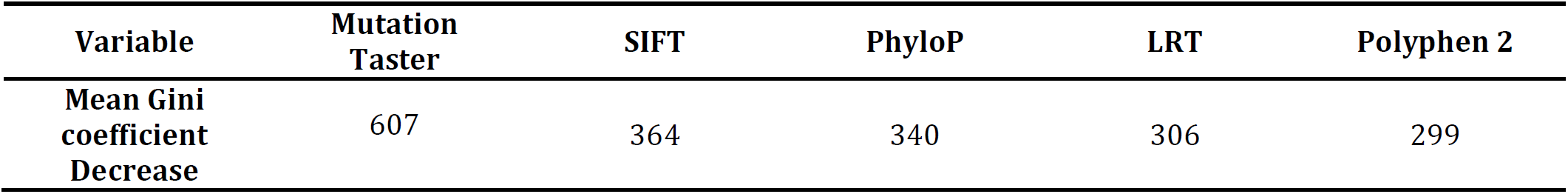
Variables importance in rfPred model

In order to address the two above named specific situations (avoiding missing a potential deleterious variant, or on the contrary, identifying only variants with a very high pathogenicity probability), we have calculated a partial AUC for very high sensitivity or specificity (between 0,9 and 1), and compared these results with the partial area index. These indices can be interpreted as the mean sensitivity (specificity) of each classifier for the studied specificities (sensitivities)^26^.

In Table 3, we can see that rfPred has a higher index than the logistic regression model on the learning dataset in both cases.

**Table 3:** Partial Area Index calculated on the learning dataset

| Models | rfPred<br>(OOB*) | Logistic<br>regression | CADD | Mutation<br>Taster | SIFT | Polyphen 2 |
| --- | --- | --- | --- | --- | --- | --- |
| **Sensitivity**<br>**between**<br>**0,9 and 1** | **0,316** | 0,285 | 0,246 | 0,241 | 0,197 | 0,220 |
| **Specificity**<br>**between**<br>**0,9 and 1** | **0,414** | 0,375 | 0,196 | 0,311 | 0,202 | 0,210 |

\*OOB = Out of bag data

### *3.2* *Model Validation on independent data*

On the four validation datasets, each time rfPred is in par with the best performing predictors (Figure 2). The rfPred AUC is 0,90 on EXOVAR dataset (vs 0,84 for Polyphen2 and 0,86 for Mutation Taster), of 0,86 on COL7A1 (vs 0,85 for Polyphen2), of 0,95 on COL4A5 (vs 0.946 for SIFT). On the ClinVar-1KG validation set, rfPred is clearly more accurate than the logistic regression framework. Although the most reliable predictor seems to be Mutation Taster on the EXOVAR dataset, on the two specific gene-based validation datasets, SIFT and Polyphen2 give better results (Table 4). The strength of rfPred is that in all these datasets, it is the only one to remain among the most accurate ones.

**Fig 2:**
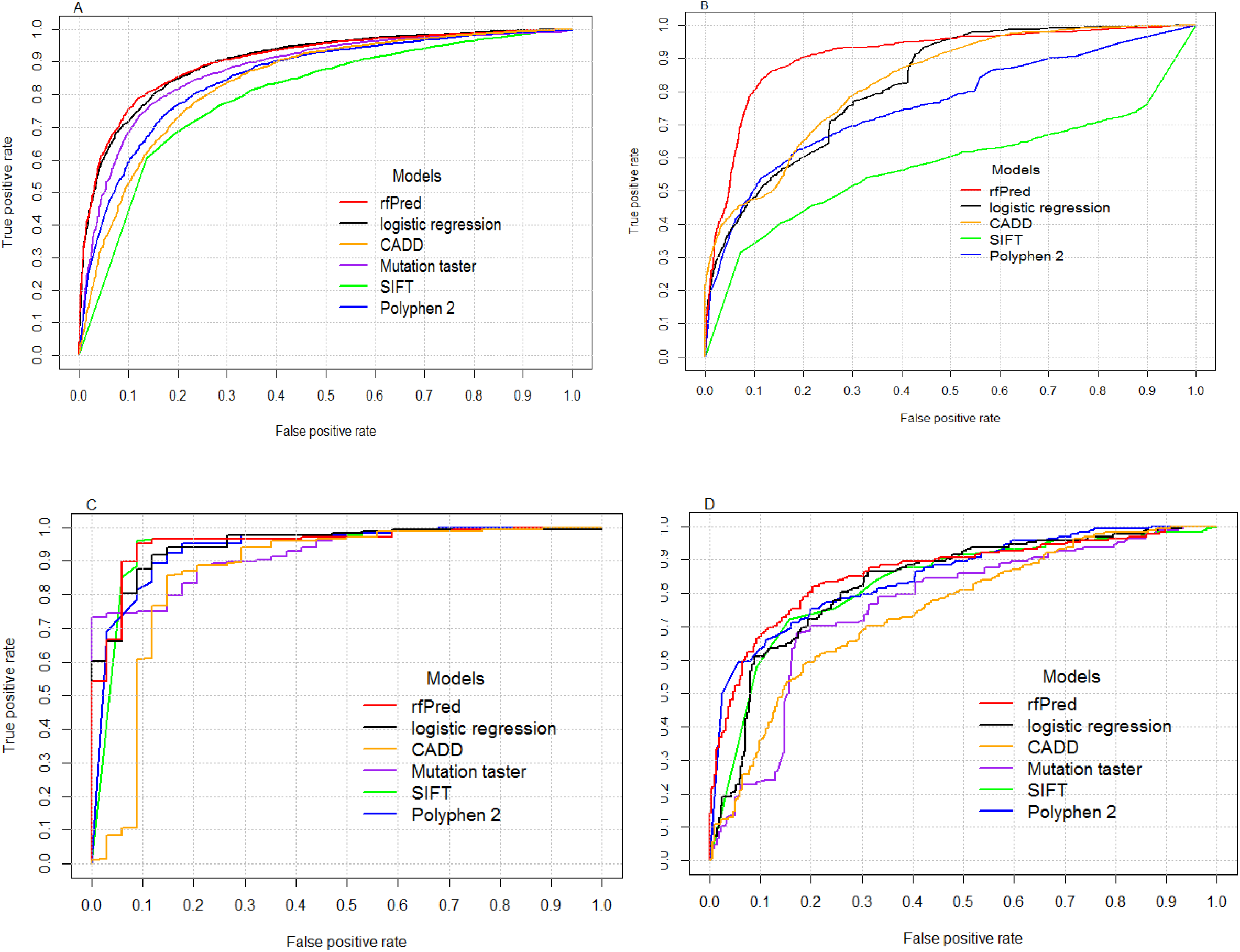
ROC curves on validation datasets for best classifiers. A: EXOVAR validation set, B: ClinVar – 1Kg validation set, C: COL4A5 validation set, D: COL7A1 validation set (LRT and PhyloP are not shown because their ROC curves are below the other ones)

**Table 4:** Models comparison on the different validation datasets

| Models | rfPred | Logistic regression | Mutation Taster | SIFT | Polyphen 2 | CADD |
| --- | --- | --- | --- | --- | --- | --- |
| Total AUC (EXOVAR) | **0,90\*** | **0,90** | 0,86 | 0,80 | 0,85 | 0,84 |
| Partial AUC Index Sensitivity between 0,9 and 1 (EXOVAR) | 0,46 | **0,50** | 0,41 | 0,25 | 0,37 | 0,39 |
| Partial AUC Index Specificity between 0,9 and 1 (EXOVAR) | **0,58** | 0,55 | 0,45 | 0,22 | 0,39 | 0,31 |
| Total AUC (ClinVar-1KG) | **0,91** | 0,82 | NA | 0,58 | 0,76 | 0,83 |
| Partial AUC Index Sensitivity between 0,9 and 1 (ClinVar-1KG) | **0,52** | 0,48 | NA | 0,02 | 0,14 | 0,41 |
| Partial AUC Index Specificity between 0,9 and 1 (ClinVar-1KG) | **0,52** | 0,35 | NA | 0,20 | 0,34 | 0,40 |
| Total AUC (COL4A5) | **0,95** | 0,95 | 0,92 | 0,95 | 0,94 | 0,87 |
| Partial AUC Index Sensitivity between 0,9 and 1 (COL4A5) | 0,72 | 0,72 | 0,53 | **0,75** | 0,70 | 0,66 |
| Partial AUC Index Specificity between 0,9 and 1 (COL4A5) | 0,73 | 0,71 | **0,74** | 0,62 | 0,63 | 0,13 |
| Total AUC (COL7A1) | **0,86** | 0,84 | 0,76 | 0,83 | 0,85 | 0,75 |
| Partial AUC Index Sensitivity between 0,9 and 1 (COL7A1) | 0,29 | **0,40** | 0,21 | 0,28 | 0,36 | 0,26 |
| Partial AUC Index Specificity between 0,9 and 1 (COL7A1) | 0,48 | 0,30 | 0,16 | 0,31 | **0,50** | 0,18 |

\*Figures in bold are corresponding to the best result of the row

If we consider now the partial AUC for high sensitivity or high specificity, the rfPred method is especially effective to detect disease causing variants with a minimum of false positives (high specificity). The logistic regression model outperforms the rfPred model for partial AUC with high sensitivity in two datasets from the four.

## 4. Discussion and Conclusion

The protein function alteration prediction of a genetic SNP variant located on a coding DNA domain is a real challenge today. Many parameters should be taken into account, more or less bounded to each other, and the existing pathogenicity prediction methods give complementary information. The added value of combining several existing predictors is well established.

Although a logistic regression framework has already been proposed and seems to improve the accuracy of the five prediction scores, we have chosen a classification model based on a machine learning approach (random Forest) so that the method can take into account the nonlinear part of the link between variant pathogenicity and their prediction scores, and can model complex interactions between them. Such interactions would be complicated to define within a classical multiplicative model framework (Supplementary Data). The resulting rfPred model seems not only to be more accurate than the logistic regression framework already proposed and the CADD score for missense variants, but is also easily available to the scientific community.

rfPred is built with 5 000 trees, which was found to be the best compromise between complexity and accuracy on our training dataset; with 10 000 trees, the classification errors do not decrease and the forest stability does not increase anymore significantly (data not shown).

Concerning the learning dataset we have deliberately chosen to work with the neutral variants which have a frequency < 1% in the 1000 Genomes Project. Indeed, more frequent variants are easily excluded from sets of possible causal variants in monogenic diseases without resorting to any statistical tool. According to us, the added-value of prediction methods for monogenic diseases appears precisely when a high number of variants remain potentially pathogenic even after filtering on the minor allele frequency in the general population. Our model is intentionally built on the more challenging dataset in term of classification; the implicit hypothesis is that the genetic variants with a frequency higher than 1% in the population are more often correctly classified as neutral by function alteration predictors. We have checked this hypothesis in comparing the rfPred scores distribution according to allele frequency of neutral variant (Supplementary data) and seen that the neutral variants with a MAF > 1% have globally lower rfPred scores than the rare neutral SNVs.

rfPred has been compared with other classification models coming from machine learning or statistical learning communities as Support Vector Machine (SVM) or boosting technique ^20^. A recent example of such a tool is the CADD method ^6^, and rfPred seems really more performant on our different datasets. CADD offers the great advantage to provide scores for all the possible Single Nucleotide Variants of the human reference genome and not only for coding variants, but in the particular field of missense single nucleotide variations, it does not seem to be the most accurate one.

It could be interesting to add other prediction scores in our composite model, in particular those which have demonstrated a very good accuracy in particular contexts, like Mutpred ^27^ or SNP&GO ^28^. The recently releases of dbNSFP v2 introduce a few others predictors which could be used as well.

Another following step could be to integrate such a model in a more complex tool like ANNOVAR or VAAST 2.0 ^29^ which allows finding candidate genes from phenotypically homogeneous exomes. VAAST 2.0 enables to join several approaches: the approach linked to the variant prioritization based on the amino-acid substitution (those which led us to develop rfPred), and the association analysis approach, based on the comparison between cases and controls. In the unified likelihood model of the tool, the variant prioritization is taken into account through an approach based on a conservation measurement PhastCons; a variant prioritization model like ours could further improve this tool, whatever its use in common diseases or in rare diseases.

Finally, to decrease the request time, rfPred scores have been pre calculated for each possible variation in the human exome and are available in tabix files downloadable at http://www.sbim.fr/rfPred. A R package downloadable on Bioconductor.org queries the data files stored directly on the server website, or locally downloaded. For a query of 200 variations scores, the request time is about 1s if the data files are stored locally on the computer (versus 6s if the data files are on the website). It is also possible to download the random forest model (in R.Data format) to compute the rfPred scores directly from the five LRT, SIFT, Polyphen2, PhyloP and MutationTaster scores.

## Acknowledgments

We thank Corinne Antignac, Alain Hovnanian and Matthias Titeux for checking our validation datasets on COL4A5 and COL7A1 genes, and Audrey Virginia Grant for her manuscript comments.

